# Mental navigation and telekinesis with a hippocampal map-based brain-machine interface

**DOI:** 10.1101/2023.04.07.536077

**Authors:** Chongxi Lai, Shinsuke Tanaka, Timothy D. Harris, Albert K. Lee

**Affiliations:** Janelia Research Campus, Howard Hughes Medical Institute, Ashburn, Virginia, USA

## Abstract

The hippocampus is critical for recollecting and imagining experiences. This is believed to involve voluntarily drawing from hippocampal memory representations of people, events, and places, including the hippocampus’ map-like representations of familiar environments. However, whether the representations in such “cognitive maps” can be volitionally and selectively accessed is unknown. We developed a brain-machine interface to test if rats could control their hippocampal activity in a flexible, goal-directed, model-based manner. We show that rats can efficiently navigate or direct objects to arbitrary goal locations within a virtual reality arena solely by activating and sustaining appropriate hippocampal representations of remote places. This should provide insight into the mechanisms underlying episodic memory recall, mental simulation/planning, and imagination, and open up possibilities for high-level neural prosthetics utilizing hippocampal representations.

## Introduction

The ability to simulate scenarios in one’s mind is a hallmark of intelligence, facilitating the evaluation of past experiences and future plans. For instance, we can imagine walking around our previous workplace, or imagine how our current workplace might function if we rearranged the furniture. Importantly, such imagination requires an internal world model that can be flexibly accessed to construct possible scenarios (Schacter et al., 2007, 2012; Hamrick, 2019)

The hippocampus is a brain region that is critical for memory and imagination (Hassabis et al., 2007a, 2007b; Addis et al., 2007; Schacter et al., 2007) and it holds a model of the environment (also called a cognitive map (Tolman, 1948; O’Keefe and Nadel, 1978)) that could potentially be mentally traversed for the purpose of recall or simulation. In particular, the hippocampus contains spatial map-like representations of previously explored environments, most prominently studied in rodents. Each environment’s representation consists of place cells—neurons that fire selectively whenever an animal moves through specific locations (called the “place fields” of those cells) in that environment (O’Keefe and Dostrovsky, 1971; O’Keefe, 1979). This results in a distinct multi-cell activity pattern at each location in the environment which, during physical navigation, can be used to decode the animal’s current location from the ongoing pattern of neural activity (Wilson and McNaughton, 1993). In contrast, a key aspect of imagination is the activation of neural representations that deviate from current sensory input. For instance, your ability to picture that walk through your old workplace while sitting in a chair at home involves activation of representations that are non-local (i.e., away from your current location). Previous work has shown brief and intermittent activation of non-local hippocampal spatial representations suggestive of the planning of specific paths within a cognitive map (Johnson and Redish, 2007; Diba and Buzsáki, 2007; Davidson et al., 2009; Karlsson and Frank, 2009; Pfeiffer and Foster, 2013; Wikenheiser and Redish, 2015; Mou et al., 2022; Widloski and Foster, 2022). However, it is unknown whether this activity is volitionally controlled, or rather reflects passive memory-related processes similar to the trajectory-like patterns observed during sleep (Louie and Wilson, 2001; Lee and Wilson, 2002) that are presumably non-volitional.

Here, we tested whether an animal can directly control its hippocampal activity according to its model of the world. We conducted this test in rodents and employed a brain-machine interface (BMI) approach because, unlike in humans, we cannot simply ask animals to imagine scenarios. Instead, with BMI methods, we could reward animals for generating neural activity resembling the simulation of specific scenarios. More precisely, we could reward them for the volitional activation of specific non-local representations from the cognitive map—a fundamental building block of scenario simulation. BMI research has a rich history of directly testing for volitional control of activity patterns of neuronal ensembles in motor cortex and related areas (Wessberg et al., 2000; Carmena et al., 2003; Musallam et al., 2004; Hochberg et al., 2006; Santhanam et al., 2006; Velliste et al., 2008; Sadtler et al., 2014; Aflalo et al., 2015; Anumanchipalli et al., 2019; Willett et al., 2021; Fetz, 2007; Lebedev and Nicolelis, 2017). In the hippocampus, it has been shown that the activity level of individual neurons (Ishikawa et al., 2014; Patel et al., 2021) or the population activity related to individual stimuli (Cerf et al., 2010) can be controlled. But, a real-time BMI that allows animals, humans or otherwise, to control their hippocampal population activity in terms of the content of their cognitive map (e.g., location representations) has never been shown. We show here that rats equipped with a real-time hippocampal BMI can navigate to goals (in our “Jumper” mental navigation task) or move external objects to goals while remaining stationary (in our “Jedi” telekinesis task) within an immersive virtual reality (VR) environment solely by controlling the activity of a population of place cells. The animals achieve this by activating and sustaining (non-local) representations of the places where they want themselves or a particular object to go. This demonstrates the kind of flexible, goal-directed manipulation of the cognitive map that could underlie recollection or simulation of richly detailed experiences.

### A hippocampal map-based BMI

Each Jumper or Jedi BMI experiment consisted of three phases (Figure 1A). In phase 1, rats ran to a succession of arbitrary locations marked by a tall, visible goal cue placed in a familiar 2-dimensional virtual arena (“Running task”). Upon reaching each cue, liquid reward was delivered, the trial ended, and the cue moved to another location for the next trial. Animals were secured in a harness and could freely rotate their body and head direction on top of a spherical treadmill (Aronov and Tank, 2014) while hippocampal CA1 neural activity was recorded (Figure 1B, Figure S1). We applied a recently developed field-programmable gate array (FPGA)-based neural signal processor to perform low-latency (1 ms) assignment of extracellular spikes (recorded from 128 channels) to a population of hippocampal units (Lai, 2020). In the Running task, treadmill movement updated the animal’s location in the virtual environment and many hippocampal units (i.e., place units) displayed spatially modulated activity (Harvey et al., 2009; Aronov and Tank, 2014; Lee et al., 2020) (Figure 1B, blue arrows) similar to that in real-world environments (O’Keefe and Dostrovsky, 1971; O’Keefe and Nadel, 1978; Wilson and McNaughton, 1993). In phase 2, the binned spike counts from the activity of these place units and the animal trajectory from the Running task were used to train a decoder (Figure 1B, green arrows) that estimates the animal’s current location from the neural data every 100 ms. As in recent motor BMI work (Anumanchipalli et al., 2019; Willett et al., 2021), we used a deep neural network for decoding (Figure S2), allowing the use of data augmentation for training—a method that improves decoder performance given limited data as well as its noise robustness. In phase 3, the treadmill was disconnected from the VR system and the animal’s ability to control its own or an object’s translational movement was limited to controlling its hippocampal activity, which was converted by the decoder into a specific location output every 100 ms (Figure 1C). Importantly, the decoder was trained to estimate the animal’s current location in the Running task only, not its location in the subsequent BMI tasks. As mentioned, during BMI periods, the animal needed to, and did, generate activity corresponding to locations away from its current location.

**Figure 1.**
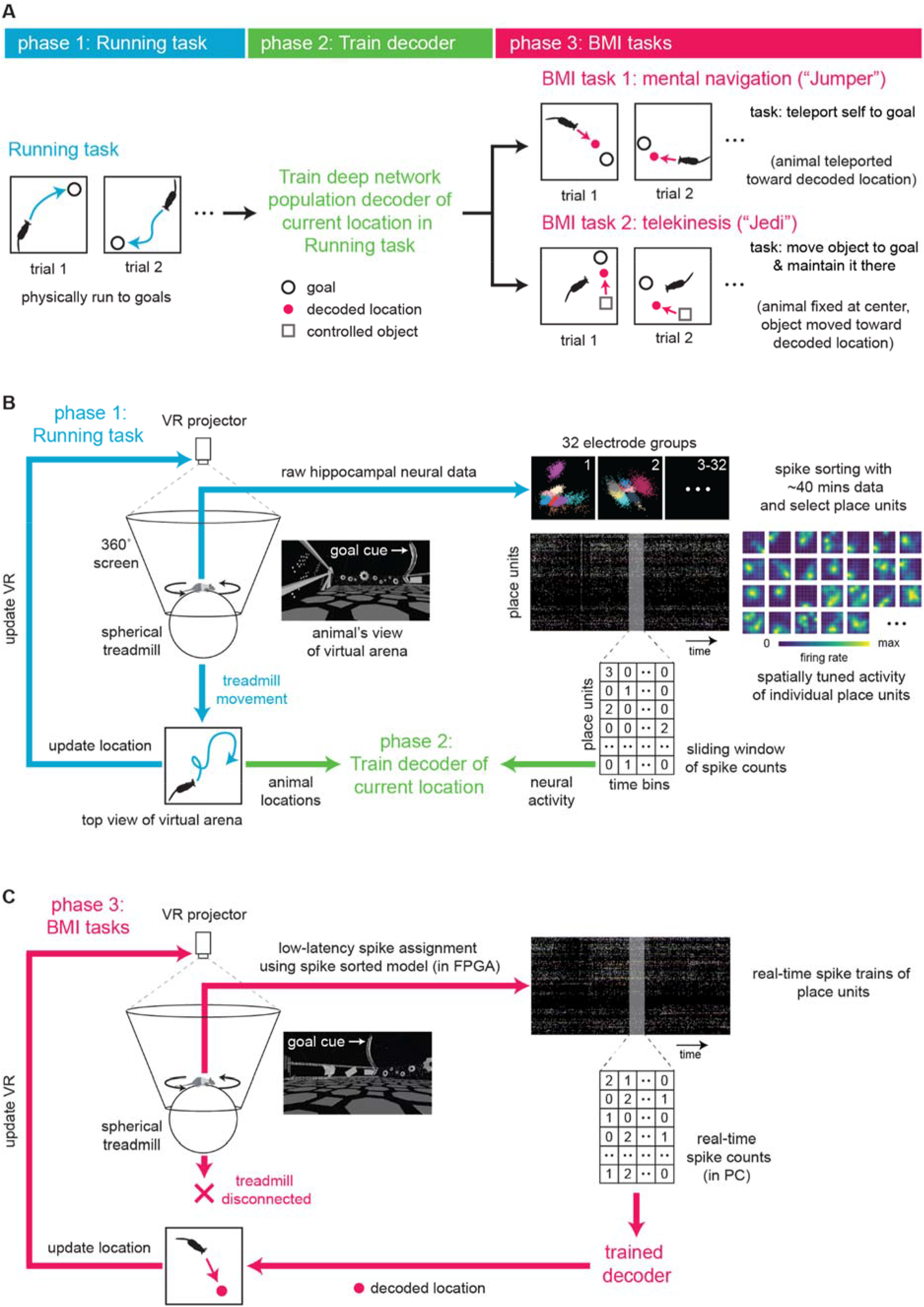
Hippocampal map-based brain-machine interface (BMI) in a virtual reality (VR) system. (**A**) Steps for performing the two different BMI experiments in this study. Rats first physically ran to a series of goals (“Running task”) while their hippocampal neural activity and (virtual) location in a square arena were recorded. This data was used to train a decoder to take neural activity as input and output the animal’s current location in the Running task. In BMI task 1 (“Jumper”), animals needed to generate neural activity that would be decoded as locations they wanted to move to so that they could reach each goal (to obtain reward). In BMI task 2 (“Jedi”), animals were fixed at the center of the virtual arena (but could rotate) and needed to generate activity corresponding to locations where they wanted an external object to move to so that the object reached the goal, then they needed to sustain that activity to maintain the object there (to maximize reward). (**B**) Schematic of VR system (left). Animal was free to rotate its body in the horizontal plane. In the Running task, animal’s location in the virtual arena environment was updated based on treadmill movement. Simultaneously recorded spiking from a population of hippocampal CA1 units expressed place fields—the basis of the cognitive map of the environment (right). Decoder was then trained using binned spiking activity and location data. (**C**) In both BMI tasks, treadmill no longer updated VR. Instead, the animal or object location was controlled solely by real-time hippocampal activity. A neural signal processor rapidly assigned activity to individual units, whose spike counts were fed into the decoder. VR projection was updated based on locations output by the decoder. In the “Jumper” (“Jedi”) task, the animal’s (object’s) virtual location was moved toward the most recent decoded locations.

### Mental navigation task

In the Jumper task, we tested whether animals could navigate to arbitrary goal locations as in the Running task, except here via BMI-based first-person teleportation. After rats performed the Running task for ∼40 min (∼120 trials) (Figure 2A,B), the data was used to train the decoder, which accurately estimated the rat’s current location in the Running task (validation set R^2^=0.78-0.88, Figure 2C). Jumper trials were identical to Running trials, except the animal’s location was updated to the BMI-decoded location (smoothed with a 3 s sliding window) (Figure 2D). If an animal did not reach the goal within 62 seconds, the trial ended and a new goal cue appeared at a random location.

**Figure 2.**
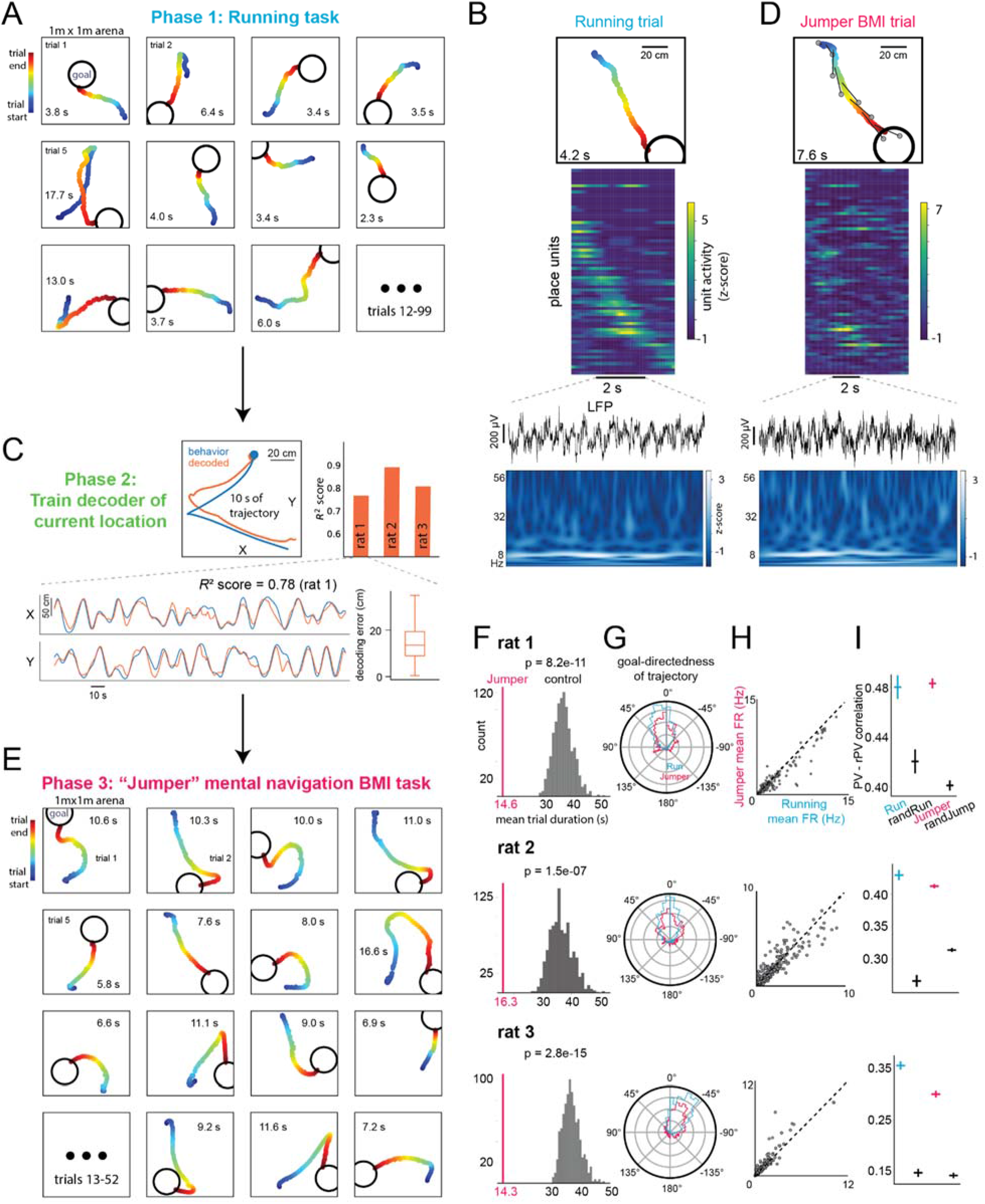
Rats can navigate to goals by controlling their hippocampal activity. In both Running and Jumper BMI tasks, animals were rewarded when they reached each goal. (**A**) Animal trajectories in virtual arena for consecutive Running task trials. Trial duration (time to reach goal) in seconds shown. (**B**) Example Running task trial. From top: trajectory, firing rate (z-scored) of individual units (units were ordered by time of peak activity), and LFP from one recording channel and corresponding wavelet spectrogram during trial. (**C**) Accuracy of trained decoder of animal’s current location for held-out Running task data. Actual and decoded trajectories during example trial (top left) and across several trials (for X and Y coordinates separately, bottom left). Median decoding error (distance between actual and decoded locations) with range and quartiles (bottom right). (**D**) Example Jumper BMI trial with similar trajectory as Running trial in (B). From top: trajectory generated by the animal controlling its hippocampal activity and the decoder output (animal is teleported toward decoded location; each gray circle represents the decoded location at the time the animal is at the corresponding point in the trajectory connected by the dark line), firing rate of individual units (using same order of units as in (B)), LFP, and spectrogram. (**E**) BMI-generated trajectories for consecutive Jumper trials. (**F**) Mean Jumper trial duration (vertical line) is significantly lower than distribution of expected mean duration for simulated trials if goals were in random locations. (**G**) Polar distribution of angle between direction of movement and direction to goal during Running and Jumper tasks. Zero corresponds to animal movement directly toward the goal center. (**H**) Mean firing rate of each unit during Jumper versus Running trials. Dotted line: y = x. (**I**) Mean correlation of instantaneous (500 ms window) population vector (PV) of activity in Running or Jumper task with mean hippocampal map PV (rPV) for the current location (in Running task), current decoded location (in Jumper task), or random location in Running (randRun) or Jumper (randJump) task. See text for details. Here and elsewhere all CIs are 95% CIs.

Rats successfully navigated by controlling their hippocampus, generating efficient paths to each goal (Figure 2E, see Figure S3-S5 for all trials of 3 rats). To check if this performance could be due to non-spatially-specific neural activity (e.g., modulating global firing rate), we randomly shuffled the spike trains across place units, ran the shuffled data through the original decoder to produce simulated trajectories, then determined how long it would have taken to reach the same sequence of goal locations as in the original experiment. Shuffled-unit mean trial durations were much longer than the actual means (*P<*10^-100^, 3 rats, 1 session each), suggesting performance depended on generating place field-related activity. Crucially, to test whether generating non-goal-directed sequences of location-specific activity (e.g., random movement within the cognitive map) could explain the performance, we randomly shuffled the goal locations in each trial while preserving the original BMI trajectories, then determined the time that would have been needed to reach the shuffled goals. Shuffled-goal mean trial durations were again much longer than actual means (*P*=2.8×10^-15^-1.5×10^-7^, Figure 2F), indicating animals’ BMI trajectories were clearly goal-directed. Goal-directedness was also apparent from the distribution of angles between the animal’s instantaneous direction of BMI-generated movement and the direction from animal current location to goal, which was concentrated around a value near 0° (Figure 2G). Thus, even though Jumper trials took longer than Running trials (mean trial duration across animals: 15.1 versus 6.9 s; note, though, that BMI decoding and smoothing added a few seconds to Jumper durations), the animals’ routes revealed effective, goal-directed, map-based BMI navigation. Furthermore, such performance was achieved without extensive BMI training (Figure 2E, Figure S3-S5 show sessions 3, 9, 2 for rats 1-3, respectively; a fourth rat failed to perform either BMI task). Initially, animals physically ran as in the Running task, but in later trials animals ran less (Figure S6).

What characteristics did the volitionally generated activity have? First, local field potential (LFP) activity during Jumper displayed prominent theta (∼5-12 Hz) band power, similar to the Running task (Figure 2B,D). This was not unexpected as the animal often, but not always, moved on the treadmill during the task. Second, population burst events (PBEs), during which brief activation of place cell representations for remote locations sometimes occur (Diba and Buzsáki, 2007; Davidson et al., 2009; Karlsson and Frank, 2009; Pfeiffer and Foster, 2013; Mou et al., 2022; Widloski and Foster, 2022; Lee and Wilson, 2002), appeared after obtaining rewards but not when approaching goals, thus did not contribute to Jumper performance. Third, mean firing rates per unit were similar between Jumper and Running tasks (Figure 2H). Lastly, we investigated the hypothesis that, to move toward a given (decoded) location in the Jumper task, animals generated a pattern of firing rates across units (i.e., a population vector, or PV) similar to the mean PV at that location over the entire Running task (called the reference PV, or rPV), i.e., similar to that location’s sample from the standard place field map across the population. Specifically, we examined the correlation between the PV generated at each moment (in every 500 ms window) during Jumper and the rPV of the decoded location at that moment. Because firing rates over such brief windows are expected to exhibit variability, as a benchmark for such variability, we first computed correlations between the 500-ms-window PVs during the Running task itself with the rPVs corresponding to the animal’s actual locations at those times and, separately, with the rPVs of random locations (Figure 2I “Run” and “randRun”, respectively). We then correlated Jumper PVs with the rPVs of the decoded locations at each moment (“Jumper”), and with rPVs of random locations (“randJump”). Jumper PVs were significantly correlated with the rPVs associated with the decoded locations (versus random locations), which is consistent with the hypothesis, and, furthermore, Jumper PV-rPV correlations were comparable to Running task PV-rPV correlations. In line with this, the example Running (Figure 2B) and Jumper (Figure 2D) trials, which happened to share similar trajectories, showed similar sequences of PVs (i.e., similar diagonals across place units).

### Telekinesis task

While episodic memories are encoded and often retrieved using a first-person perspective, individuals also can imagine scenarios from a third-person perspective, with other animate and inanimate players taking part. Furthermore, imagination often involves holding a single thought in mind for extended periods. Therefore, our second BMI task, Jedi, tested whether animals could—while remaining in place—use the same map of the arena to control the location of a virtual object, guide it to the goal cue location, and maintain it nearby. Jumper and Jedi thus employed different forms of feedback: self-location and the location of an object, respectively. After the same Running task and decoder training phases as in Jumper, the animals in Jedi were fixed (but could freely turn) at the arena’s center and the object’s location was updated to the BMI-decoded location (with a 2 s smoothing window). In each trial, the goal cue remained in place, providing reward as long as the object touched it. After 3 min or the rat received 0.5 mL reward, whichever came first, the goal cue appeared at a distant random location for the next trial.

Rats demonstrated a clear ability to activate and sustain a remote location’s representation around the goal until the trial ended, and then shift attention to the newly generated goal (Figure 3A,B, Figure S7). Performance was measured using the mean distance (over time) between the decoded locations and goals. Shuffling spike trains across units yielded much greater mean distances than the actual means (*P*=2.2×10^-5^-2.6×10^-3^, 3 rats, 1 session each). To assess the goal-directedness of BMI-generated activity, we shuffled the goal locations while preserving the locations output by the decoder. The decoded (and controlled object’s) location was far more concentrated around the actual remote goal cue than shuffled goal locations (*P*=1.8×10^-22^-5.1×10^-10^, Figure 3C, Figure S7), indicating clear goal-directed control of activity. Again, such performance occurred without extensive training (Figure 3A shows sessions 5, 6, and 3 for animals 1-3, respectively). Task performance was not dependent on population burst events (PBEs), as there was no change in performance when all activity in PBEs was eliminated and the decoder was re-run post-hoc (Figure S7). Mean firing rates per unit were correlated across Jedi and Running tasks, but lower in Jedi (Figure 3D)—consistent with decreased physical movement in Jedi (Figure S8). As in Jumper, PVs generated during Jedi were significantly correlated with the corresponding rPVs (Figure 3E). However, unlike Jumper, correlation scores were clearly lower in Jedi than the Running task, consistent with noisier generation of non-local representations in Jedi—possibly because animals often sustained remote representations for extended periods.

**Figure 3.**
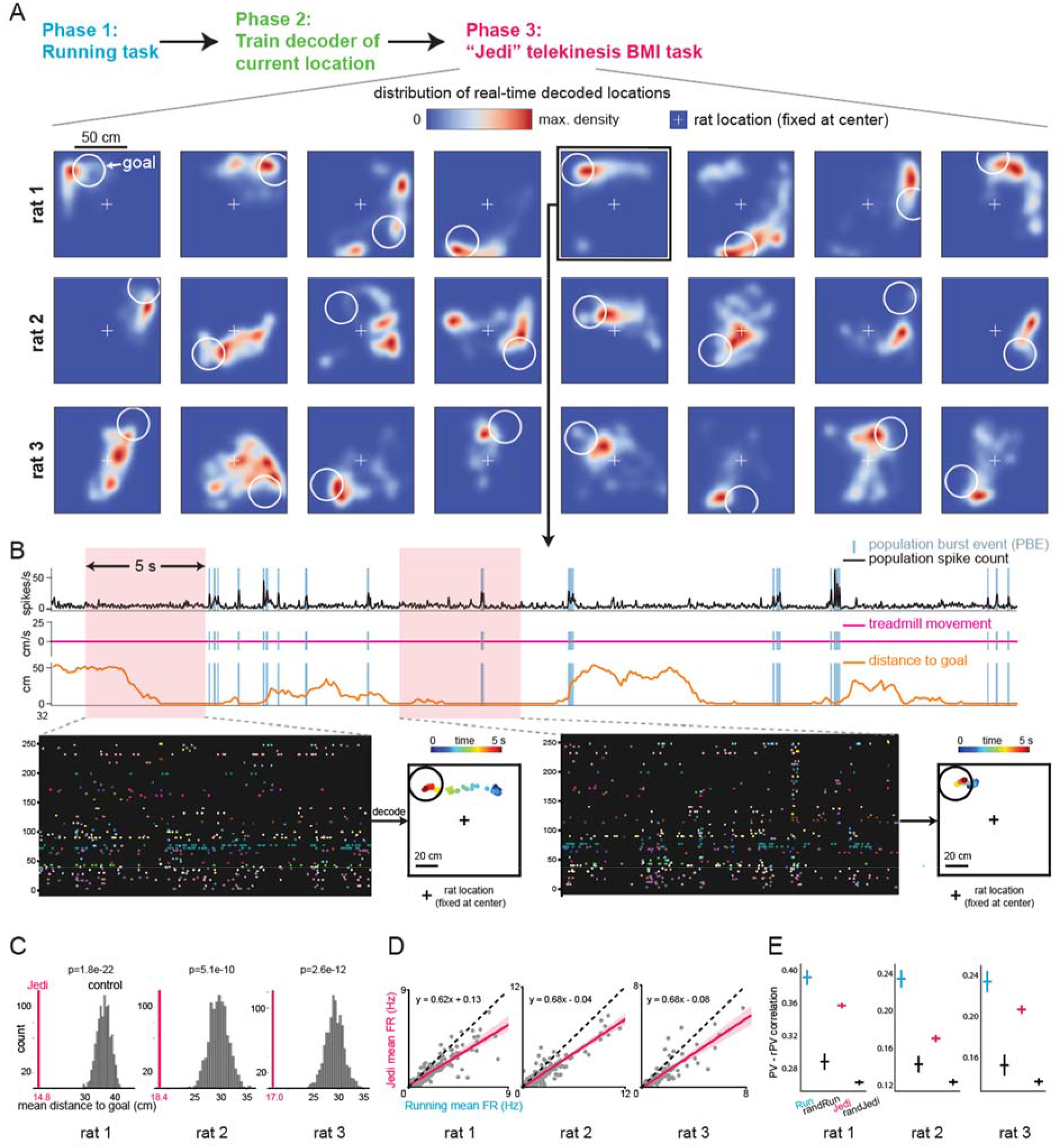
Rats can move objects to remote goal locations and maintain them there by controlling their hippocampal activity. In the Jedi BMI task, trials did not end when the external, controlled object first reached the goal; instead, animals were rewarded as long as the object was in the goal region (white circle), for up to 3 min per trial. (**A**) Distribution of real-time decoded locations (output every 100 ms) generated by the animal controlling its hippocampal activity across 8 consecutive Jedi BMI trials for rats 1-3. Panels show decoded locations during each trial (up to 3 min, Figure S7); animals did not move the treadmill the majority (71.4-85.3%) of the time shown (Figure S8) (see text and methods for details). The external, controlled object (which was visible for rats 1-2, invisible for rat 3) was moved toward the decoded location (Figure S7 shows that the distribution of object locations was essentially the same as the distribution of decoded locations). Animal was always fixed at center of virtual arena, but could rotate its body and generally turned toward each goal. (**B**) A 40-second-long period during an example trial. From top: Summed activity across all units with population burst events (PBEs) identified, treadmill speed (here, 0 for the entire period), distance of object from goal (0 means inside goal region) and close-ups of two 5-second periods (left: as animal moves object to goal; right: as animal maintains object at goal; points in arena represent sequence of decoded locations) with spike trains of units. (**C**) Mean distance of decoded location from goal across all trials (vertical line) is significantly lower than mean distance expected for randomized goal locations. (**D**) Analogous to Figure 2H. (**E**) Analogous to Figure 2I.

## Discussion

Previous BMI research has yielded major advances in the control of robotic arms, computer cursors, and other devices by activity from primary motor, premotor, and posterior parietal cortex. Advances have included increases in the speed of controlled output, use of different decoding outputs such as movement parameters or motor intentions, and investigation into the intrinsic neural constraints on what activity patterns a subject can learn to control (Wessberg et al., 2000; Carmena et al., 2003; Musallam et al., 2004; Hochberg et al., 2006; Santhanam et al., 2006; Velliste et al., 2008; Sadtler et al., 2014; Aflalo et al., 2015; Anumanchipalli et al., 2019; Willett et al., 2021; Fetz, 2007; Lebedev and Nicolelis, 2017). Importantly, the hippocampal cognitive map has a code that represents space in terms of absolute location in the external environment versus location relative to (e.g., in front or to the right or left of) the animal (O’Keefe and Dostrovsky, 1971; O’Keefe and Nadel, 1978; Wilson and McNaughton, 1993), and it was unknown whether a subject could control a BMI based on this code. Here, for the first time, we have demonstrated a hippocampal map-based BMI in which the subject is able to control its location or that of other objects by activating location representations in terms of absolute space, independent of where the animal currently is. That is, even though animals generally (but not always) turned their body toward the goal (partly to see where it is), the activity that needed to be generated differed depending on the location of the goal with respect to the environment. The relatively small amount of training needed for the animals to perform our BMI tasks is in line with our use of such a biomimetic decoder (Shenoy and Carmena, 2014; Lebedev and Nicolelis, 2017), i.e., one based on the neural code that the subject naturally employs.

Previous work in humans revealed that imagining or recalling objects or video clips is accompanied by hippocampal activity in individual neurons similar to that when viewing the original stimuli (Kreiman et al., 2000; Gelbard-Sagiv et al., 2008). This suggests that the mechanisms allowing animals to selectively activate their non-local hippocampal spatial representations as shown here could also underlie our ability to actively recall or imagine experiences in other places. The ability of rodents to perform these BMI tasks should thus allow imagination, as well as the voluntary recall of memory, to be investigated using the range of tools available for this model system. More generally, the neural processes engaged here could underlie our capacity to perform “mental time travel” (also called autonoetic consciousness), i.e., travel back in time by re-experiencing richly detailed episodic memories and forward in time by generating possible future scenarios (Tulving, 1985). Mental time travel depends critically on the hippocampus (Hassabis et al., 2007a, 2007b; Addis et al., 2007; Andelman et al., 2010; Race et al., 2011; Bartsch et al., 2011) and enables subjects to internally simulate new experiences according to their world model. This can aid decision-making and facilitate learning in complex situations where trial and error is expensive, as shown using artificial agents (Weber et al., 2017; Ha and Schmidhuber, 2018; Hafner et al., 2019; Hamrick, 2019).

Along these lines, we found that our animals, surprisingly, were able to control their hippocampal map-based activity on a timescale of seconds, corresponding to the speed and duration at which humans relive past events or imagine new scenarios. Navigational trajectories each lasted ∼10 s, and a virtual object could be held at a remote location for tens of seconds. This contrasts with the previously described fast (∼100 ms) sequences of non-local hippocampal activity in awake rodents thought to be associated with planning (Johnson and Redish, 2007; Pfeiffer and Foster, 2013), and which were not responsible for the performance in our BMI tasks (population burst event analysis). While it is not yet known whether the content of these fast sequences can be under an animal’s volitional control, these events could represent a brief consideration of alternatives for making a quick decision and be distinct from the more comprehensive mental simulations of possible scenarios that take seconds.

Beyond service in decision-making, the ability to control the contents of the hippocampal spatial and episodic memory system could help explain the richness of our inner lives. Finally, the ability to control hippocampal activity to guide oneself or objects to intended locations—and do so with high signal-to-noise read out using our decoder—could lead to new BMI applications for restoring or enhancing function based on realizing a subject’s high-level intentions with respect to their internal world models.

## Acknowledgments

We thank M. Bolstad for work on the virtual reality software; W. Bishop, S. Romani, and X. Zhao for valuable discussions regarding the study; S. Lindo, R. Gattoni, and other members of the Janelia Vivarium team, and B. Lustig, A. Sohn, S. Sawtelle, I. Negrashov, B. Foster, J. Osborne, J. Arnold, R. Rogers, and J. Cohen for technical assistance; and J. Dudman, S. Romani, A. Hantman, T. Wang, and J. Colonell for valuable comments on the manuscript. This work was supported by the Howard Hughes Medical Institute.

## Author contributions

Conceptualization and task design: C.L.; BMI and task software and BMI hardware: C.L.; behavioral and recording hardware: S.T.; experiments: S.T., C.L.; data analysis: C.L., A.K.L.; writing: C.L., A.K.L., S.T.; supervision: A.K.L., T.D.H.

## Methods

### Subjects

The subjects were adult male Long-Evans rats, weighing ∼350-450 g at the time of surgery. Animals were individually housed in home cages fitted with custom-made running wheels throughout training and after surgery on a 12 h light / 12 h dark schedule. Animals were water-restricted to provide motivation to perform the Running and BMI tasks, during which liquid reward could be obtained. All procedures were performed according to the Janelia Research Campus Institutional Animal Care and Use Committee guidelines on animal welfare.

### Virtual reality system

#### Virtual environment software and hardware

Our custom virtual reality (VR) software was developed as part of Janelia’s virtual reality software platform (Jovian). Our software suite, named ‘MouseoVeR’, was written in C++ and built from a number of open-source software components (Boost, Bullet, osgBullet, osgWorks, OpenSceneGraph, Collada, OpenGL, and Qt) (Cohen et al., 2017). Virtual arena environments were created using the open-source animation software Blender (www.blender.org) and rendered by MouseoVeR. Blender environments were rendered using six virtual camera objects located at a single point in space to cover all the directions of a cube. For display, the six images were converted to an annulus shape to be projected from above onto a custom-made screen (80% polyester, 20% spandex, First Response Custom Sewing, Inc.). The screen was shaped like an inverted truncated cone stretched between two aluminum rings (top and bottom ring diameters: 122 cm and 63.5 cm, respectively, height: 100 cm). The final image encompassed a viewing angle of 50° above and 30° below the horizontal eye level. The top ring was mounted on an aluminum frame. The top of the screen was covered by a horizontal sheet of transparent acrylic (122 × 122 × 1.2 cm). A projector (InFocus 5312a) with short-throw lens (InFocus LENS-060) was mounted horizontally on the ceiling, and images (1600 × 1200 pixels) of a virtual environment were reflected at a right angle off a round mirror (18-inch diameter, McMaster-Carr) onto the screen through the acrylic sheet. The cone-shaped screen surrounded a spherical treadmill, which consisted of a large, hollow, lightweight polystyrene sphere (24 in diameter, 350 g total when the 2 separate hemispheres were glued together, Foam Mart ball modified by WeCutFoam) resting on a bed of seven individually air-cushioned ping-pong balls (30 psi) arranged around the lower half of the sphere by an acrylic frame (https://www.janelia.org/open-science/large-spherical-treadmill-rodents (Cohen et al., 2017)). To prevent chewing of the sphere by the rats, the sphere was covered with packing tape (Scotch). Rotation of the sphere around its vertical axis was prevented by four “yaw blockers” (small, vertically oriented rubber wheels with custom made attachments) separated by 90° around the sphere’s equator, thus the animals could only change direction by rotating themselves on the treadmill. To track the motion of the treadmill, two cameras separated by 90° were positioned at the equator and focused on 4 mm^2^ regions under infrared illumination (modified from FlyFizz, (Seelig et al., 2010)). The cameras captured 30 × 30 images of the treadmill surface at 4 kHz and the motion of the treadmill was computed from the accumulated differences in the images over time. In each iteration of the rendering loop, MouseoVeR communicates with the treadmill’s data server to retrieve the updated motion values since the last request. For calibration, we created mappings between 180° rotations of the treadmill along each of the two relevant directions in real-world space and the data server’s coordinate space. Body orientation in the horizontal plane was detected by a rotational encoder (TRD-MX1000AD, Koyo) that was calibrated by a photointerruptor (EE-SX672-WR, Omron) and a Teensy 3.1 microcontroller. The animal was held in place at the apex of the treadmill by a body harness, consisting of a jacket for the forelimbs (Harvard Apparatus), two spandex belts supporting the lower body (16 × 1.5 cm), and a neoprene backbone (10.5 × 5.5 cm, 70A hardness). A 3D-printed hook on this backbone was used to fasten the animal to an aluminum arm containing two hinges, which permitted a small amount of up-and-down body movement during walking. The arm was attached to a bearing (X-contact, 3.500 × 4.000 × 0.25, Swerve Drive Specials) on the bottom acrylic plate of a 128-channel motorized commutator (Saturn, Neuralynx), which allowed the rat to freely turn its body in any horizontal direction on the treadmill (i.e., 360° free rotation with no restriction on the number of net turns it could accumulate in either the clockwise or counterclockwise direction). Another 3D-printed arm that also rotated with the animal was attached to hold a lick port in front of the animal. To not block the animal’s forward view, this arm had an open window (16 × 12 cm) in front of the animal. On the top of the commutator, a liquid rotary joint (Doric) was mounted, and its rotation was linked to the commutator so that the water reward supply line could go through the commutator and freely rotate with it. The back end of the water supply line was connected to two solenoid valves (EV-2-12, Clippard) and a syringe pump (PHD2000, Harvard Apparatus) to control the amount and timing of water reward. In addition, a 50 mL reservoir and a small solenoid valve (SV-2C-12-3-V, Clippard) were mounted onto the aluminum arm so that sweetened water (Kool-Aid) could also be provided, if desired. Whenever reward was delivered, a buzzer beeped (SunFounder, 2300 ± 300 Hz). On the acrylic frame holding the spherical treadmill, four nozzles (Eppendorf 1000 µL tips connected to stainless steel tubing) were attached 30 cm away from the animal to supply airpuffs (30 psi). Airpuffs were triggered manually when animals did unfavorable behaviors, such as chewing the equipment or sitting still for too long (e.g., during Running task training). The screen, projection, yaw restriction, commutator, and body harness systems were modeled after (Aronov and Tank, 2014).

#### VR game engine for task implementation

The Running, Jumper, and Jedi tasks were all written in Python using our VR game engine called Playground. Playground (https://github.com/chongxi/playground) is a Python-based software system that allows for the creation, execution, and control of complex behavioral tasks in VR environments. At the core of Playground is a Finite State Machine (FSM) framework used to define the task’s logic and rules. For instance, when a trial starts, the task state is set to “trial started”. A cue is then generated at a random location. Once the animal moves close enough to the goal, the cue triggers a transition in the task state from “trial started” to “reward cue touched”, then to “reward delivering”, and finally to “trial finished”. Each state transition can lead to a specific outcome. The state transition and associated outcome is fully defined in the FSM. This FSM is fully customizable and simple to prototype with using a few lines of Python code, allowing for the creation of a wide range of tasks, from simple go/no-go paradigms to more complex, multi-state tasks. Playground also includes a user-friendly visualization module that enables researchers to intuitively track task states, animal behavior inside a 3D-modeled environment, and electrophysiological data, such as spike trains and waveform features, simultaneously in real time. With this fast 3D visualization, researchers can observe and understand the relationship between behavior, neural activity, and task states for rapid BMI task prototyping. Playground also integrates with the Jovian VR platform, which is used in this study to render the 360° immersive virtual environment. The combination of Playground and Jovian software allows for the precise control and manipulation of the virtual environment, including the ability to teleport objects and change their properties in real time according to either animal behavior or BMI output. The code for all of the tasks that were used in this study can be found on the GitHub page of the Playground project (https://github.com/chongxi/playground/blob/master/playground/base/task/task.py).

### Brain-machine interface system

#### Overview

Our brain-machine interface system performs real-time analysis and converts the place unit activity of a population of CA1 units into an estimate of the animal’s current location in the Running task (and the “desired” location in the BMI tasks). It consists of a Field Programmable Gate Array (FPGA)-based neural signal processor (NSP) and a deep neural network (DNN)-based decoder, the latter of which resides on a personal computer (PC). The NSP is connected to up to 5 32-channel headstages (RHD2132, Intan Technologies) that amplify and sample neural signals from up to 160 channels (128 channels were used in this experiment) at a rate of 25 kHz per channel. The NSP communicates with the PC through Python application programming interfaces (APIs), which allow the NSP to retrieve the parameters of a spike sorting model from the PC. The FPGA uses these parameters to classify spikes as belonging to individual units in 1 ms, and sends the assigned spike-id’s (which identify the units) along with spike timestamps to the PC through a low-latency interface for real-time decoding. The FPGA-based NSP is described in more detail below in this section, and the DNN-based decoder in the BMI task section.

Our BMI system was integrated with the VR systems. During the pre-BMI Running task, band-pass filtered raw recordings, together with online-detected spike waveforms, the waveform features, and their electrode origins, as well as the animal’s location and body orientation at each moment, were collected and stored. After conducting semi-automated spike sorting, place units were selected based on their activity level and spatial information for use in training the decoder. This decoder was trained to estimate the animal’s current location in the Running task based on recent neural activity from the population of place units. This decoder was then deployed in real time for BMI tasks. In the BMI tasks, the online-decoded location was transmitted to the VR game engine, enabling updates of the task state, and from the game engine to MouseoVeR for VR rendering.

#### FPGA-based NSP for on-chip spike assignment

The real-time FPGA-based neural signal processor for classifying spikes into their source unit is described in detail in (Lai, 2020). Briefly, the 128 channels of raw data that were amplified and sampled at 25 kHz per channel (32 bits per sample) by the headstages were input into the FPGA (KC705, Xilinx). Inside the FPGA, (1) each channel was band-pass filtered between 500-3000 Hz using a custom pseudo-linear phase FIR filter, (2) common noise across channels was removed by reference subtraction, (3) the data were split into 32 independent 4-channel electrode groups (2 per 8-site shank), (4) within each electrode group, spikes were detected using an amplitude threshold (see below) and 19 points (0.76 ms) per channel around each spike’s peak were assigned as the (4 × 19 dimensional) waveform for that spike, (5) principal components of each spike waveform were extracted as features, based on principal component analysis (PCA) that had been applied to all the spike waveforms from the corresponding electrode group acquired during the training period (here the Running task), (6) each spike was classified as coming from a given unit with respect to unit clusters that had been defined from spike sorting the training period data. Importantly, the PCA and classification model, used for identifying the unit origin of each spike, were determined offline by analyzing and curating the training data on a PC. The resulting PCA transformation matrix and classification model for each electrode group were manually inspected, then transferred to the FPGA for fast, online processing. In the FPGA, the PCA matrix converts online-detected spike waveforms into waveform features, which are then classified by comparing them to pre-determined reference features for each unit. The time elapsed from spike detection to the completion of spike classification is a deterministic latency of 1 ms for each spike, regardless of the number of spikes or units due to the real-time processing power of FPGAs (Lai, 2020).

### Behavioral training

#### Acclimation to equipment

To acclimate the rats to wearing the harness and behaving on the treadmill, a series of steps were followed. First, the rats were subjected to water restriction for 1-2 weeks. Then, they began a habituation process in which they wore the jacket for 10-15 minutes each day, then the jacket plus belts, until they became comfortable with them. This typically took about 10 days. Then they were placed in the full harness on the treadmill for a few days with ample water provided from the lick port. During this time, they were encouraged to start obtaining water regularly from the lick port. This acclimation process helped the rats become accustomed to the harness and treadmill, which allowed for goal-directed running and BMI task behavior.

#### Training using the Running task

After acclimation, animals were exposed to a cue-rich VR environment (1 × 1 m square arena with 20 cm high walls) with proximal cues (on the floor and walls) and distal cues (around the arena above the walls). A tall, thin, spiral pillar (of maximum extent 20 × 20 × 80 cm) was used as the goal cue for all tasks. To increase its visual salience, the goal cue moved up and down at a frequency of approximately 1 Hz. During the initial training phase, the goal cue was placed close to the animal. Whenever the animal reached within 20 cm of the goal cue (center-to-center distance, called the “goal radius”), it was rewarded with 20-40 µL of water or 40-50 µL of sweetened water, and the goal cue was moved to a new location. This training phase served to help the rat understand that touching the goal cue leads to reward. Once the animals demonstrated consistent navigation to the goal cue, the Running task was introduced, in which the goal cue was placed at a random location at least 50 cm away from the location of the last reward, and the goal radius was decreased from 20 to 15 cm (or sometimes 10 cm). The same VR environment was used for all of the training and experiments in order to allow the animal to form a well-learned map of the arena for the BMI experiments.

### Surgery and electrode targeting

After an animal’s performance in the Running task improved to the point of reaching the goal and obtaining reward ∼200 or more times per 90 min training session, the animal was anesthetized with isoflurane and mounted in a stereotaxic frame for chronic implantation of electrodes. Two craniotomies were made, one over the CA1 field of the dorsal hippocampus of each hemisphere (centered at AP −3.7 mm, ML 2.8 mm). The dura was removed, and a 64-channel silicon probe consisting of 8 shanks with 8 recording sites each (Buzsaki64-H64LP_30mm, Neuronexus) was inserted into each hemisphere at an initial depth of ∼900 µm. Each silicon probe was mounted on a shuttle drive (Nano Drive, Ronal Tool Company), which was fixed to the skull using OptiBond Universal (Kerr), Charisma A1 (Kulzer), dental cement (Jet Acrylic, Lang), and Calibra Universal (Dentsply Sirona). The probes were each connected to two of the 32-channel headstages (RHD2132, Intan Technologies), and the probes and headstages were surrounded by custom-made 3D-printed protective shells (the headstages remained with the implant when the animal was in its home cage). After a week of recovery, the electrodes were gradually adjusted over several weeks until they reached the CA1 pyramidal cell layer. Electrophysiological features such as the amplitude and polarity of sharp waves and the amplitude of spikes, recorded during each day’s ∼90 min training session in the Running task, were assessed visually to guide adjustment to the CA1 cell layer. After performing the BMI experiments, animals were anesthetized with isoflurane, small electrolytic lesions were made by passing anodal current (30 µA, ∼10 s) through one electrode site per hemisphere, then animals received an overdose of ketamine and xylazine and were perfused transcardially with saline followed by 4% formaldehyde. Brains were coronally sectioned (50 µm thick) and placed on slides with mounting media containing DAPI (Vectashield) to verify recording locations.

### Brain-machine interface tasks

#### Running task before either BMI task

After CA1 place unit activity was observed during a Running task training session (from offline sorting of data and analysis of spatially tuned firing, as in Figure 1B), we started BMI experiments for that animal. On a given day, we first recorded neural activity for ∼10 s to set the spike threshold (mean minus 4.5 to 5 × the standard deviation of the activity (Quiroga, 2012)). Then, animals were required to perform ∼40 min of the Running task (∼120 trials) while neural activity was recorded. Animals had to get to within 15 cm (or, in some sessions, 10 cm) of the center of the goal cue to get reward. Afterwards, animals were temporarily returned to their home cage, which was placed in the same room as the VR system.

#### Semi-automatic spike sorting

For the spikes that were detected online using threshold crossing per channel during the Running task, the four waveforms per spike from that electrode group (set of 4 adjacent electrodes) were saved, and these saved spike waveforms were used for offline semi-automatic spike sorting (which was done separately per electrode group). The spike waveforms of an electrode group were PCA-transformed with 4 principal components kept, then a Dirichlet process Gaussian mixture model was applied (using the scikit-learn package in Python) to cluster the spike waveform features of each electrode group into 15-20 clusters (i.e., units). Using this “over-split” spike sorting model (i.e., one place cell might be split across more than one unit) reduces the amount of time needed to curate the unit clusters. This is important because a shorter curation time results in a smaller gap between the end of the training period (i.e., the Running task) and the start of the BMI task, thus reducing the potential impact of electrode drift. Additionally, using an over-split spike sorting model, which is related to clusterless decoding (Kloosterman et al., 2014; Deng et al., 2015), leads to the same or better population decoding performance compared to single-unit spike sorting by maximizing the amount of information obtained from each electrode group. Finally, rapid manual curation was conducted using the interactive 3D visualization software of our BMI system, during which we separated noisy or unstable units from stable units by visual inspection, and a spike-to-unit classification model was built from this manually curated clustering result. We removed the noisy or unstable units from further processing (i.e., building the location decoder) while keeping these clusters for the online spike assignment to absorb noisy spikes. In this manner, many “noise” spikes were accurately labeled as belonging to noisy clusters and therefore did not contribute to either offline or online location decoding. However, some of the noise spikes (particularly when small electrode drifts occur) can invade the boundary of the well-isolated clusters. Such noise is likely inevitable, but we handle noise by explicitly training the location decoder to be less sensitive to noise in general (see below).

#### Spatial tuning (place fields), spatial information, and selection of place units

To determine the spatial tuning of a unit, a two-dimensional histogram of the unit’s spiking locations in the 1 × 1 m arena (binned into 4 × 4 cm spatial bins) was first generated. The histogram was then normalized by the total duration the animal spent in each spatial bin. The resulting firing rate map was subsequently smoothed using a Gaussian filter with a standard deviation of 8 cm. Only periods of time when the rat’s speed was >5 cm/s were used to calculate place fields. The collection of firing rate maps across the units represents a sample from the hippocampus’ spatial map of the environment (cognitive map). Given the firing rate map *f* of a unit, the information rate *I* (bits/spike) of a unit was calculated as follows (Skaggs et al., 1993):

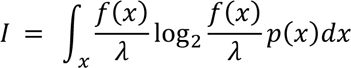

where *λ* is the mean firing rate of the unit and *p*(*x*) is the probability that the rat is present in the spatial bin *x*.

Place units were selected from the pool of all stable units to train the decoder. Selection criteria included the peak rate of the unit’s spatial firing rate map and the unit’s information rate, both of which had to exceed certain thresholds, and which varied across animals (default parameters were set to exclude units with <0.5 Hz peak firing rate, spatial information <0.1 bits/spike, or <0.1 Hz or >4 Hz mean firing rate, the last of which should generally exclude interneurons, leaving units from pyramidal cells). The number of (oversplit) stable units and, of these, the number of stable place units satisfying the above criteria, were: for Jumper (157 and 61 units for rat 1, 373 and 122 for rat 2, 233 and 159 for rat 3, respectively), and for Jedi (253 and 110 units for rat 1, 288 and 124 for rat 2, 138 and 94 for rat 3). Note that only the place units were used for training the decoder, real-time decoding in BMI tasks, re-decoding analysis, and PV analysis. We used all stable units for the firing rate analysis in Figure2H and Figure3D as well as for population burst event (PBE) detection.

#### Deep neural network decoder to estimate animal’s current location in the Running task

To estimate the current location of an animal, we trained a 16-layer DNN (Figure S2) using the neural activity and locations during animal movement (counting only those periods when the speed, smoothed with a zero-lag 2 s boxcar window, was >5 cm/s) in the pre-BMI Running task. The DNN was trained to minimize the Euclidian distance between its estimate of the current location and the animal’s actual current location (smoothed with a zero-lag 3 s boxcar window) every 100 ms. The input to the DNN consisted of the last 5 seconds of CA1 population activity, discretized into 50 100 ms time bins (note that in one session, the rat 3 Jumper session, the last 1.5 s of CA1 population activity was used instead as the input). Each bin contains the spike counts produced by N place units. To stabilize the spiking noise variance, the elements of the N × 50-dimensional (or N × 15-dimensional, when using the last 1.5 s of activity) spike count matrix were square-rooted, in accordance with methods proposed previously (Kihlberg et al., 1972; Yu et al., 2009). The DNN transformed the N × 50-dimensional spike count matrix into a single 256-dimensional vector and passed it through all the internal layers until reaching the final linear layer, which output a 2-dimensional vector representing the x and y coordinates of the estimated animal location. Inspired by recent research that employs periodic nonlinear activation function (Sitzmann et al., 2020), we designed a new network backbone that employs a sinusoidal activation function in place of the commonly used rectified linear unit (ReLU). We named this block Sine Net (Figure S2). We observed that a stack of this network structure using skip connections between each instance of the Sine Net significantly accelerated the training on some of our data in comparison to other nonlinear activation functions.

#### Data augmentation to improve decoder robustness

To improve the robustness of our DNN location estimation in the presence of input noise (which could come from online spike sorting, motion artifact, electrode drift, physiological state changes, and other sources), we employed data augmentation (Willett et al., 2021) that adds independent and identically distributed (IID) noise to the training data. This noise was generated using a Bernoulli random variable and a uniform random variable (between 0 and a maximum value). The Bernoulli variable introduced a probability of 0.5 that the spike count of each place unit at each time bin would be changed (either increased or decreased) independently. The uniform variable determined the magnitude of this change, which was sampled independently for each unit and each bin. To ensure that the input remained valid, we set the spike count to zero whenever it became negative after the change. During training, the DNN was exposed to various levels of noise by using the absolute value of a sinusoid as the maximum magnitude of the noise across epochs (Figure S2F). This data augmentation procedure allowed the network to learn to ignore independent noise across units (i.e., “off-manifold” activity). By training the DNN model to produce similar outputs in the presence of various amounts of noise, we improved its robustness and generalization performance. (Note that data augmentation is only used during training but not the evaluation of decoder performance.)

For the Jedi task (rats 1-3) and one session of the Jumper task (rat 3), factor analysis (FA)-based reconstruction of the spike count matrix was conducted before adding the independent noise. FA is a dimensionality reduction technique widely used in BMI (Yu et al., 2009; Santhanam et al., 2009; Sadtler et al., 2014) that models shared variability and maps the population neural activity onto its intrinsic manifold. Akin to PCA, FA is used when the underlying structure of the data is believed to be a linear combination of uncorrelated latent variables, called factors, but, importantly, FA separates the neural population activity into two components: shared-variability that is generated by a set of low-dimensional factors (on-manifold activity) and independent noise that can be different across units (off-manifold activity). This explicit separation provides an additional source for data augmentation during training which could further improve our decoder robustness. To achieve this, latent factors (40-dimensional) were used to reconstruct raw spiking activity by removing off-manifold activity, ideally resulting in only on-manifold activity, then independent noise was added across units to create an additional set of training data. Because FA reconstruction (using the first 40 factors) reduces off-manifold activity in the raw input, this further encourages the DNN to rely more on the “on-manifold” activity (Sadtler et al., 2014) for location estimation. Our FA-based data augmentation was implemented in Python using scikit-learn and PyTorch.

#### Evaluation of decoder performance

We used the last 70% of the unit data from the Running task to train the current location decoder and tested the performance of the decoder on the first 30% of the data. This procedure took ∼10-20 min. After training, the decoder was evaluated on the test set using the coefficient of determination (i.e., R^2^ score). To compute the R^2^ score, the decoder output was first smoothed with a zero-lag 3 s boxcar window (as was the actual current location in the test set). (Note that the decoder output was not smoothed during DNN training.) The mean R^2^ score was calculated by taking the mean of the R^2^ scores for the x and y axes separately and was used as the final performance score of each individual session (Figure 2C). A higher value of R^2^ indicates more accurate decoding.

If the performance of the decoder on the test data had a mean R^2^ score >∼0.8, we proceeded with one of the BMI tasks: we uploaded the spike sorting model parameters to the FPGA, placed the animal back into the VR system, had the animal run 20 trials (i.e., 20 different goal cue locations) of the Running task to visually verify the performance of the decoder, then started the BMI task (Jumper or Jedi). Note that on a given day, after the Running task, usually one session, but sometimes two sessions, of the Jumper or Jedi task were run (but the Jumper and Jedi tasks were generally not run on the same day). Rats 2 and 3 were first exposed to the Jumper task, while rat 1 was first exposed to the Jedi task.

#### Deployment of DNN decoder to control VR in real time

Spike counts from each unit were recorded and binned in real time into 100 ms bins based on their FPGA timestamps. The square roots of these binned spike counts, organized into a matrix with dimensions N × 50 (or N × 15), were sent to the trained decoder every 100 ms for location estimation. The real-time spike binning and decoding were implemented in Python and communicated to the VR game/task engine Playground using the PyTorch multiprocessing module, with each process running on a dedicated CPU and sharing data through shared memory. The parameters of the DNN were fixed after training, except for the running mean and variance in the batch normalization layer, which were updated every 60 s during the BMI tasks. The entire process of binning and decoding consistently took less than 50 ms on a PC (Dell Precision Tower 7910 with 40 CPU cores and 128 GB RAM), well within the 100 ms update interval. To smooth out any sudden jumps of the output to be rendered during BMI tasks, a 3- or 2-second moving average (i.e., an average of the last 30 or 20 outputs of the decoder) was applied to the output of the DNN to determine the updated location of the animal (in Jumper) or controlled object (in Jedi), respectively. This moving average introduced a delay of a few seconds in the BMI trajectory of animal or object relative to the current decoded locations. For both tasks, the new location that was determined every 100 ms and a command string were sent to our VR game engine to update both the task states and the VR rendering. The VR update typically took 1-2 rendering frames to complete. Note that the VR projector runs at 60 Hz and Playground (the VR engine) sends updates to the MouseoVeR VR system at 20 Hz, while the decoder updates the animal or object location at 10 Hz.

#### Mental navigation task (“Jumper”)

In the Jumper task, which is a BMI version of the Running task, the animal’s location in the virtual arena was decoupled from the treadmill movement; instead, animals were teleported to the average decoded location of the latest 3 s of decoder outputs (i.e., 30 consecutive decoding windows) every 100 ms, as described above. Whenever the rat reached within 15 cm (or, in some sessions, 20 cm) of the center of the goal cue, the current trial ended, the cue disappeared, a reward (20-40 µL of water or 40-50 µL of sweetened water) was delivered, and the goal cue reappeared at a new, random location at least 50 cm away from the animal’s location, as in the Running task. The goal radius was fixed once the session started and remained the same for all trials in that session. If the animal failed to reach a given goal cue within 62 s, the trial was also considered ended, and the location of the goal cue was changed, at which point the next trial began.

#### Telekinesis task (“Jedi”)

In the Jedi task, animals were virtually fixed at the center of the arena, the goal cue was placed at least 30 cm away from the center, and a visible (or invisible) controlled object (a rectangular object 20 × 20 cm wide and 30 cm tall with large, open sides) was teleported every 100 ms to the average decoded location of the latest 2 s of decoder outputs (i.e., 20 consecutive decoding windows). Whenever the controlled object remained within 15 (or, in some sessions, 20 cm) of the goal cue, reward could be triggered. Each reward pulse lasted 10 ms (0.3-0.4 µL of water or 0.2-0.3 µL of sweetened water) for each 100 ms decoding step, but a refractory period was implemented 0.3 s after reward delivery was triggered (i.e., after every 3 pulses). The duration of the refractory period followed a uniform distribution between 0 and 2 s. As in Jumper, the goal radius was fixed once the session started and remained the same for all trials in that session. For each trial, the location of the goal cue was fixed at a given location for 3 min or until the animal received at least 0.5 mL of reward, whichever occurred first. After that, the trial ended, and the goal cue’s location was changed while the controlled object’s location remained the same, thus starting the next trial, at which point the rat needed to control the external object to enter the new goal region.

#### Rationale for Jumper and Jedi task design

The Jumper task was designed to assess whether animals can mentally navigate to arbitrary goal locations in a goal-directed and model-based manner. To demonstrate this, animals should navigate toward each goal without searching other locations in the arena. Otherwise, animals would likely need to search the arena by producing random activity (random in the sense that it is not model-based, directed activity) until the decoded location reaches the current goal. Since the goal region only accounts for less than 7.1% or 12.6% of the arena for a 15 cm or 20 cm goal radius, respectively, not being able to use a world model (i.e., map of the environment) should result in BMI trajectories that search through many regions of the arena before finally reaching the goal and, thus, longer trial durations. Therefore, we believe performance in the Jumper BMI task is sufficient to determine whether animals can mentally navigate using their learned world model in a goal-directed manner.

However, the Jumper task, which was designed as a first-person perspective game, makes it difficult to answer three additional and potentially important questions: first, whether animals can perform mental navigation while remaining stationary; second, whether animals can hold remote locations in mind for extended periods of time, similar to what occurs in human mental time travel; third, whether animals can control an external object using the same world model (here, the same spatial map) as when controlling their own location during mental navigation. The continuous updating of the arena view and associated optical flow in the Jumper task encourages animals to move. It also encourages animals to quickly activate representations of successive locations on the way to the goal. In addition, because the trial ends immediately when the goal is reached, this tends to limit opportunities to observe prolonged periods of activation of a given remote location.

Therefore, we designed the Jedi task as a third-person perspective game that differs from Jumper in two key ways. First, the view of the arena is stable since the animal’s location is fixed at the center of the arena. Such a fixed first-person perspective view will not produce forward-moving optical flow, and thus may reduce animal movement. Second, a Jedi trial does not end when the goal cue is reached, which thus allows for a much longer trial duration (here, up to 3 min) compared to Jumper trials (which typically last ∼15 s) and allows for repeated and/or continuous activation of the representation of the goal location. These two features enabled us to observe whether animals can activate and maintain non-local representations around the goal region while remaining stationary for extended periods of time. Indeed, rats can be stationary for tens of seconds (i.e., as long as several Jumper trials) while performing the Jedi task, and they can also hold the decoded location (again, without physical movement) near or within the goal region for a tens of seconds as well, as shown in the example in Figure 3B.

### Post-experiment data analysis

Data were analyzed in Anaconda Python 3.8. All confidence intervals are 95% CIs. Two types of analyses are used in this study: “in-experiment analysis” and “post-experiment analysis”. In-experiment analysis was conducted after the Running task then applied to the BMI experiment, and is described above (e.g., semi-automatic spike sorting, selection of place units, training the DNN decoder, evaluating the decoder performance). Post-experiment analysis was conducted after the experiment and is described below.

#### The animal’s BMI performance in the Jumper task

The performance in the Jumper task was assessed by the duration of trials compared to a randomized goal control (Figure 2F). In an individual Jumper trial, a new goal cue is randomly generated away from the previous one when the last trial has finished (i.e., the animal reaches the previous goal and reward is delivered). The new goal cue may be placed behind the rat or within the rat’s visual field. When the cue is generated behind the rat, the animal cannot at first see where the cue is and typically initiates the trial by turning its body to search for the cue after consuming the previous reward (during reward consumption the rat is stationary). The time between finishing the reward and turning the body to search for the goal cue is variable, as this behavior is self-initiated by the rat. For example, the rat can groom between trials. To accurately quantify the trial duration, the trial start is defined as the moment the animal starts to engage in the task during that trial. Specifically, when the cue is presented behind the animal, the trial start is defined as the moment the rat’s body orientation changes by more than 12 degrees per second, as the rat needs to turn its body to search for the cue. When the cue is presented within the visual field of the rat and the rat does not turn to approach the goal cue, the trial start is defined as three seconds after the last trial ended, during which time the rat typically finishes consuming the reward. The trial duration is defined as the time elapsed from the trial start to the moment the goal cue is reached by the rat. To determine if the animal’s behavior in Jumper was more goal-directed than chance would predict, we performed 1000 independent shuffled goal simulations and compared the mean trial duration of each simulation with the actual mean trial duration. In each simulation, the goal locations were randomly shuffled for each trial while the animal’s BMI trajectories remained unchanged from the original data. A simulated trial ends when the BMI trajectory reaches the randomly shuffled goal cue (i.e., within the goal radius) and the simulation ends when the BMI trajectory of the entire session has been completely used. We then compared the mean trial duration of the actual Jumper session (containing ∼50 trials) to the distribution of simulated mean trial durations. The p-value was estimated from the Z score of the actual mean duration compared to the shuffle distribution. A low p-value from this test (along with a shorter actual mean trial duration) indicates that the animal’s behavior in the Jumper task was significantly more goal-directed than chance would predict.

As a further measure of goal-directed control, we computed the angle between the instantaneous direction of movement along the BMI trajectory and the direction from the current location to the center of the goal. The trajectory along each trial was divided into ∼20 samples equally spaced in time during the trial. For each sample, the angle between the direction from the last movement step and the direction from the current location to the goal’s center was computed. The distribution of angles from all samples of all trials were plotted in polar coordinates. An analogous distribution was computed for all Running trials. Both Jumper and Running task distributions were concentrated around a value that was near 0 degrees (where 0 represents movement directly toward the center of the goal). Interestingly, when the peak of the Running task angle distribution for an animal was shifted slightly away from 0, that animal’s Jumper task angle distribution was similarly shifted (Figure 2G), which indicates that animals preferred to approach goals in a similar manner when running and during BMI behavior.

In addition, performance was assessed by the duration of trials compared to a shuffled unit control. In each of 200 independent shuffles, the spike trains of each unit were randomly assigned (once across all trials) to different units, the shuffled data was input into the original decoder, and the resulting BMI trajectory determined. Then the simulated durations of each trial were computed based on how this trajectory reached the original sequence of goals locations. We then compared the mean trial duration of the actual Jumper session to the distribution of simulated mean trial durations, and estimated a p-value based on the Z score. A low p-value (along with a shorter actual mean trial duration) indicates that the animal’s performance in the Jumper task depended on the specific activity of place units, as opposed to non-spatially-specific modulation of the aggregate activity across units.

#### The animal’s BMI performance in Jedi tasks

The performance in the Jedi task was evaluated using the Euclidean distance between the controlled object and the goal cue compared to a randomized goal control (Figure 3C). The trial duration is not used as a performance metric in Jedi as a trial does not end when the object reaches the goal the first time, and a trial can last up to 3 min even if the controlled object is statistically close to (but not always within) the goal region. Like in Jumper tasks, periods of non-task-engagement, such as grooming or “trying to run out of the arena” (i.e., periods of constant running into a wall), are excluded from the analysis. Additionally, because the Jedi task was intended to evaluate the animal’s ability to control a remote object while remaining as stationary as possible, low angular movement was utilized as an indicator of task engagement, and therefore periods during which the rat’s body orientation changed by more than 12 degrees per second were excluded from the analysis. Note that, after removing these excluded periods, the majority of the time the animal did not move the treadmill (Figure S8). After excluding such non-engagement periods, a 2D histogram (with 2 × 2 cm spatial bins, smoothed by a 4 cm Gaussian kernel) of the decoded location distribution or controlled object location distribution (which were virtually identical, see Figure S7) during each trial was plotted as a visual assessment of how close the object was to the goal. To determine whether the decoded locations were closer to the goal locations than chance would predict, we compared decoded-location-to-goal distances to those from 1000 independent simulated control experiments. In each simulation, we first randomly selected a starting frame within the first 7 seconds of the Jedi session. Then, we subsampled the animal’s decoded location every 7 s (i.e., every 70th frame). This 7 s interval ensured that the decoded locations were independent of each other because the decoding window was 5 s and we used an additional 2 s moving average for determining the location of the controlled object. For each subsampled decoded location, a random goal location was generated at least 30 cm from the center of the arena (this is because in the actual Jedi experiment each goal was at least 30 cm from the center). The distance between the decoded location and the goal location was determined and 15 cm was subtracted and, if the value was less than 0, the result was set to 0. This was done because it reflects the true distance from the decoded location to the goal as far as receiving reward is concerned by accounting for the goal radius. Then, for each simulated session (1000 simulations) and the actual Jedi session, a mean distance to goal averaged across all the of the subsampled decoded locations was calculated. The actual session’s mean distance to goal was then compared to the distribution of 1000 simulated mean distance to goal values. The p-value was estimated from the Z score of the actual mean distance compared to the distribution of simulated mean distances. A low p-value (along with a smaller actual mean distance) indicates that the animal’s behavior in the Jedi task was significantly more goal-directed than chance would predict.

In addition, performance was assessed by the object-goal distance compared to a shuffled unit control. In each of 200 independent shuffles, the spike trains of each unit were randomly assigned (once across all trials) to different units, the shuffled data was input into the original decoder, and the resulting BMI object locations determined. Then the simulated object-goal distances were computed between these object locations and the original sequence of goal locations. We then compared the mean distance for the actual Jedi session to the distribution of simulated mean distances and estimated a p-value based on the Z score.

#### Re-decoding of Jedi experiment with population bursts events (PBEs) excluded

To investigate the contribution of population burst events (PBEs) to the high performance of controlling remote objects in the Jedi experiments, we detected and excluded these PBEs then applied the location decoder to the remaining neural activity and assessed the task performance post-hoc. A histogram of all place units was created with 10 ms bins and smoothed using a Gaussian kernel (with a 10 ms standard deviation). Segments where the peak of the smoothed histogram exceeded the mean plus 1.8 standard deviations were identified as candidate PBEs, with the start and end boundaries determined when the smoothed histogram crossed its mean value. Candidate PBEs of less than or equal to 10 ms duration were not counted as PBEs to reduce false positives, and the rest were counted as PBEs. Finally, the spike counts of all units in any 100 ms bin that overlapped with any PBE were set to zero, the original decoders used in each Jedi experiment were then used to re-decode the data, and the spatial distribution of resulting controlled object locations with respect to the goal location in each trial was computed (Figure S7).

#### Population vector analysis

The fact that our deep neural network (DNN) performed well during the Jumper and Jedi tasks suggests that animals can voluntarily generate goal-directed, non-local spatial activity. In addition, it suggests that the DNN generalizes well from the training set (physical navigation activity during the Running task) to new unseen data (mental navigation and telekinesis activity during the BMI tasks). To explore what information our DNN might be relying on for this generalization, we considered the evidence for one reasonable hypothesis, which is that the DNN is using spiking patterns similar to the population vectors of the spatial firing rate maps for the units at each location. We first binned the arena into 4 × 4 cm spatial bins, as we did when calculating place fields. For each spatial bin, we constructed a reference population vector (rPV), which is a vector of the averaged firing rates of all units at that spatial bin during the Running task. Next, for each moment (500 ms) during the Jumper and Jedi tasks, we calculated the Pearson correlation coefficient between the current PV (i.e., the vector of average firing rates of the units during that 500 ms) and the rPV of the DNN’s current decoded location. For comparison, we also calculated the Pearson correlation coefficient between the PV at each (500 ms) moment in the Running task and the rPV of the animal’s current location in the environment, which provides a benchmark for the “maximum” PV-rPV correlation values that can be expected when taking into account the natural variability of neural activity (especially over a timescale of 500 ms); importantly, note that the PVs (samples from the spatial map) from the Running task were the same data that went into computing the rPVs (the spatial map, i.e., the mean of those PV samples). We then compared the distribution of correlation coefficients across all moments during a session (Jumper, Jedi, or Running task) to the distribution of coefficients between the PV at each moment and the rPVs of random locations. Our results show that instantaneous PVs corresponding to each decoded location in Jumper or Jedi displayed significant (versus random levels of) similarity to the rPVs at those locations (Figure 2I and Figure 3E). This is consistent with the animal generating PVs for each location that are similar to the rPVs for those locations in the place field map, and with the DNN extracting that PV information to estimate the animal’s current location in the Running task and generated location in the BMI tasks.

#### Local field potential (LFP) and wavelet analysis

In our real-time FPGA-based NSP system, only band-pass filtered raw data within the frequency range of 500-3000 Hz was recorded. To recover the full-band raw data, a Wiener deconvolution, which was validated on both ground-truth and simulated data, was applied to the saved filtered raw data. Subsequently, one electrode in the pyramidal layer where many spikes from place units were detected was selected for LFP analysis per session. The LFP signal was obtained by applying a low-pass Butterworth filter (cutoff frequency of 400 Hz, order of 5) to the reconstructed raw data from the selected channel (sampled at 25000 Hz). LFP signals were then downsampled from 25000 Hz to 1000 Hz. Wavelet spectrograms were then obtained using complex Morlet wavelets (σ = 5) across a range of frequencies from 0-60 Hz for specific time segments. The magnitude of the spectrogram was Z-scored.

**Figure S1.**
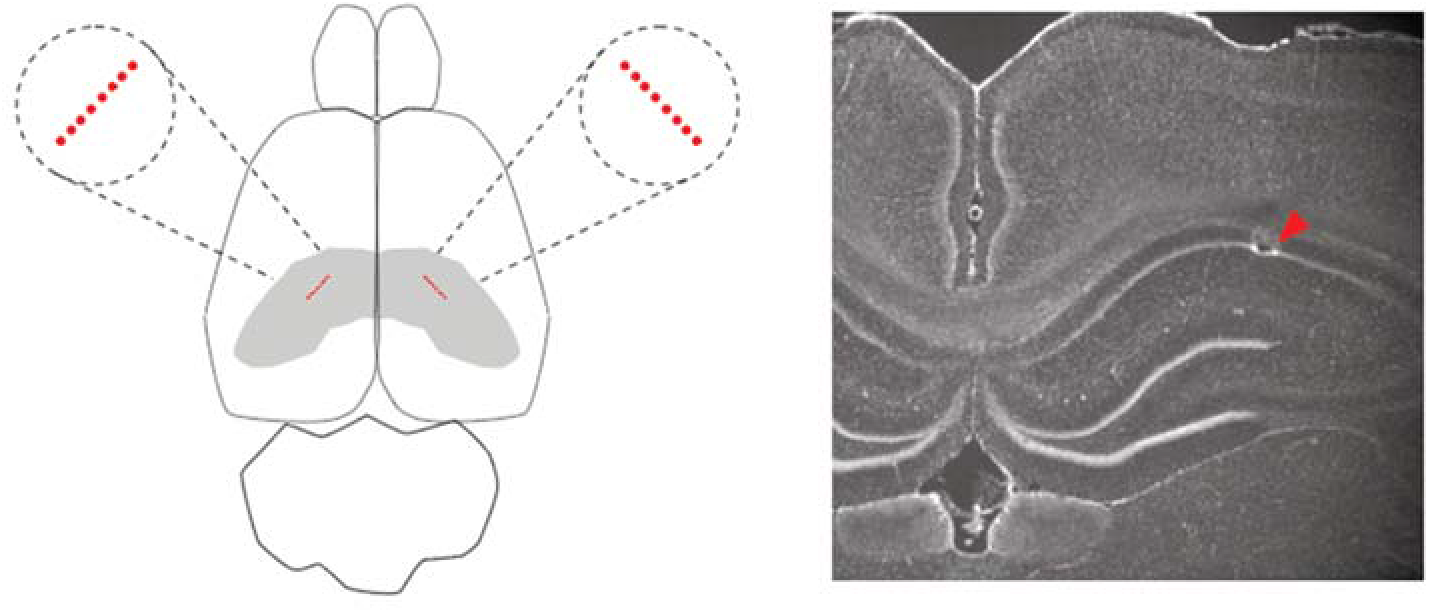
Electrode locations. Schematic of location of silicon probe shanks (each red point represents a shank) with respect to top view of brain and hippocampus (shaded) (left). Example of electrode recording site location in hippocampal CA1 pyramidal cell layer (right).

**Figure S2.**
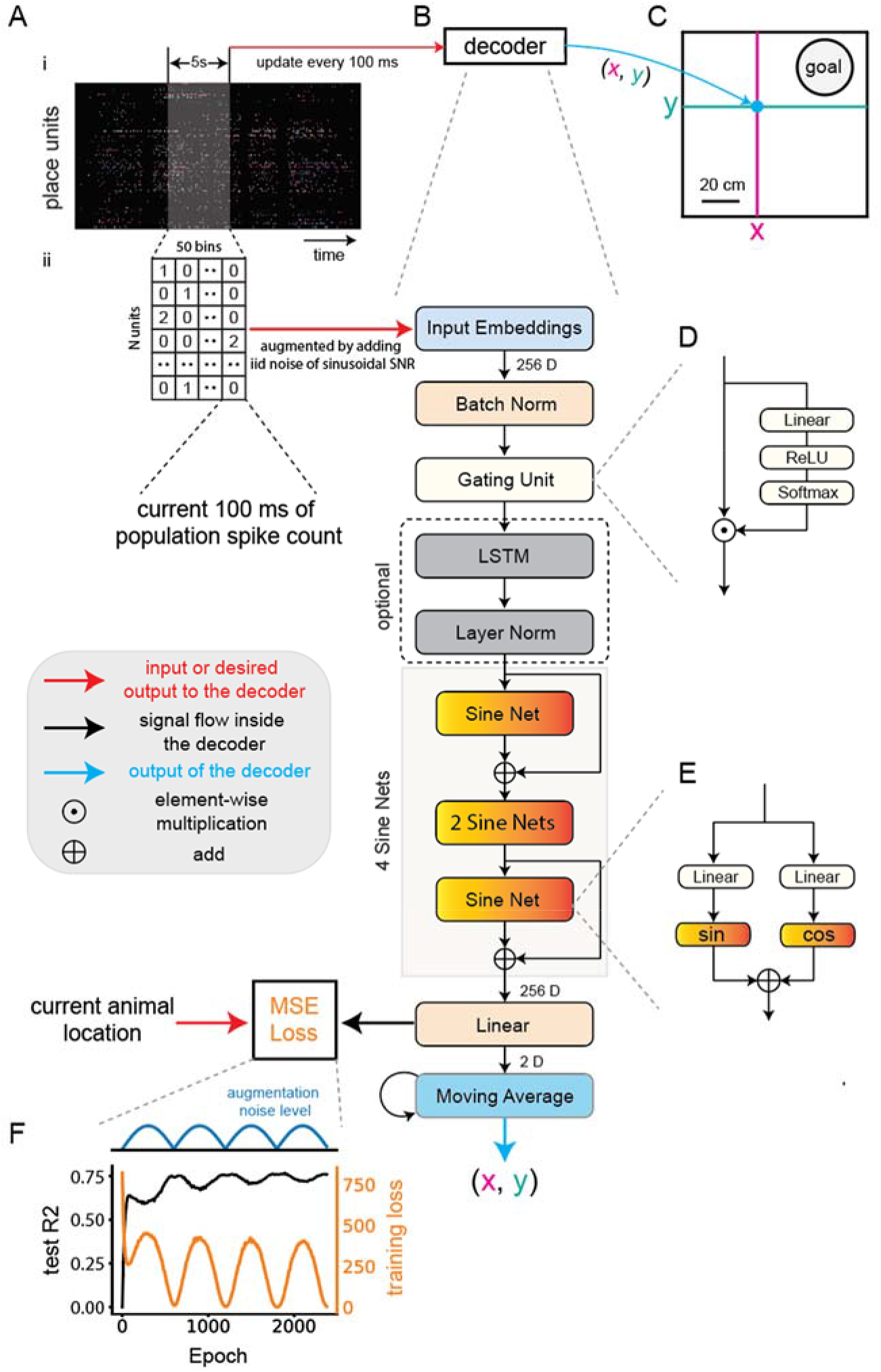
Architecture and training of deep network that accurately decodes the animal’s current location during the Running task and is used to translate hippocampal activity into control signals during the BMI tasks. (**A**) Population activity (spike counts per unit) in windows of 1.5 or 5 s were binned in 100 ms bins and input to the decoder. Windows were advanced in steps of 100 ms to give a series of inputs. (**B**) The layers and operations of the deep net decoder. (**C**) The outputs of the decoder were an x and y location value. (**D**) and (**E**) show details of gating unit and sine net layers. (**F**) The decoder was trained by minimizing the error between the current location of the animal (smoothed with a zero-lag 3 s boxcar window) and the decoder output. The training data was augmented by adding noise, which varied as shown as a sinusoidal function of training epoch. See methods for details.

**Figure S3.**
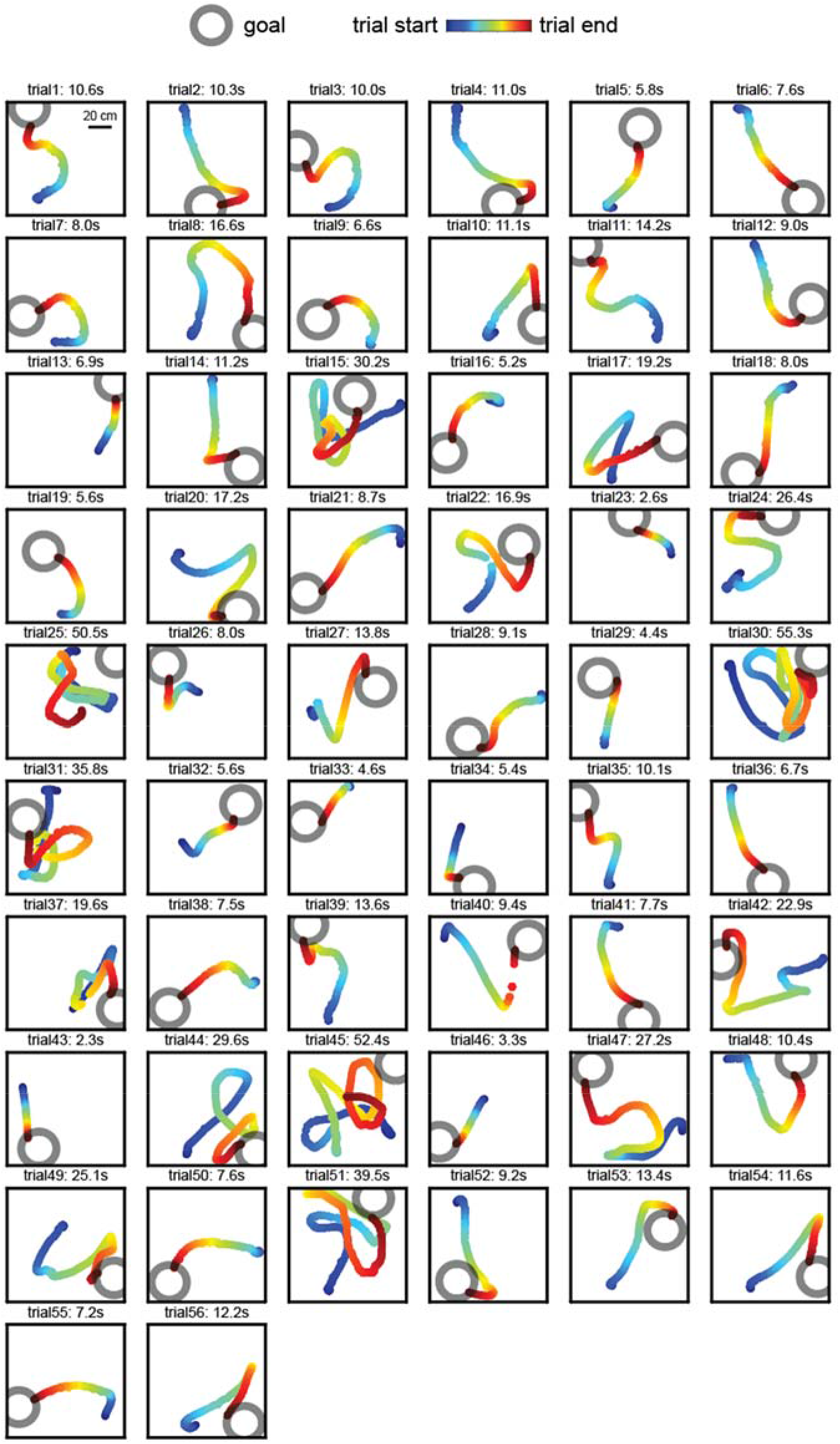
All trials of the Jumper BMI experiment for rat 1. Note trials 1-12 and 53-55 are also shown in Figure 2E.

**Figure S4.**
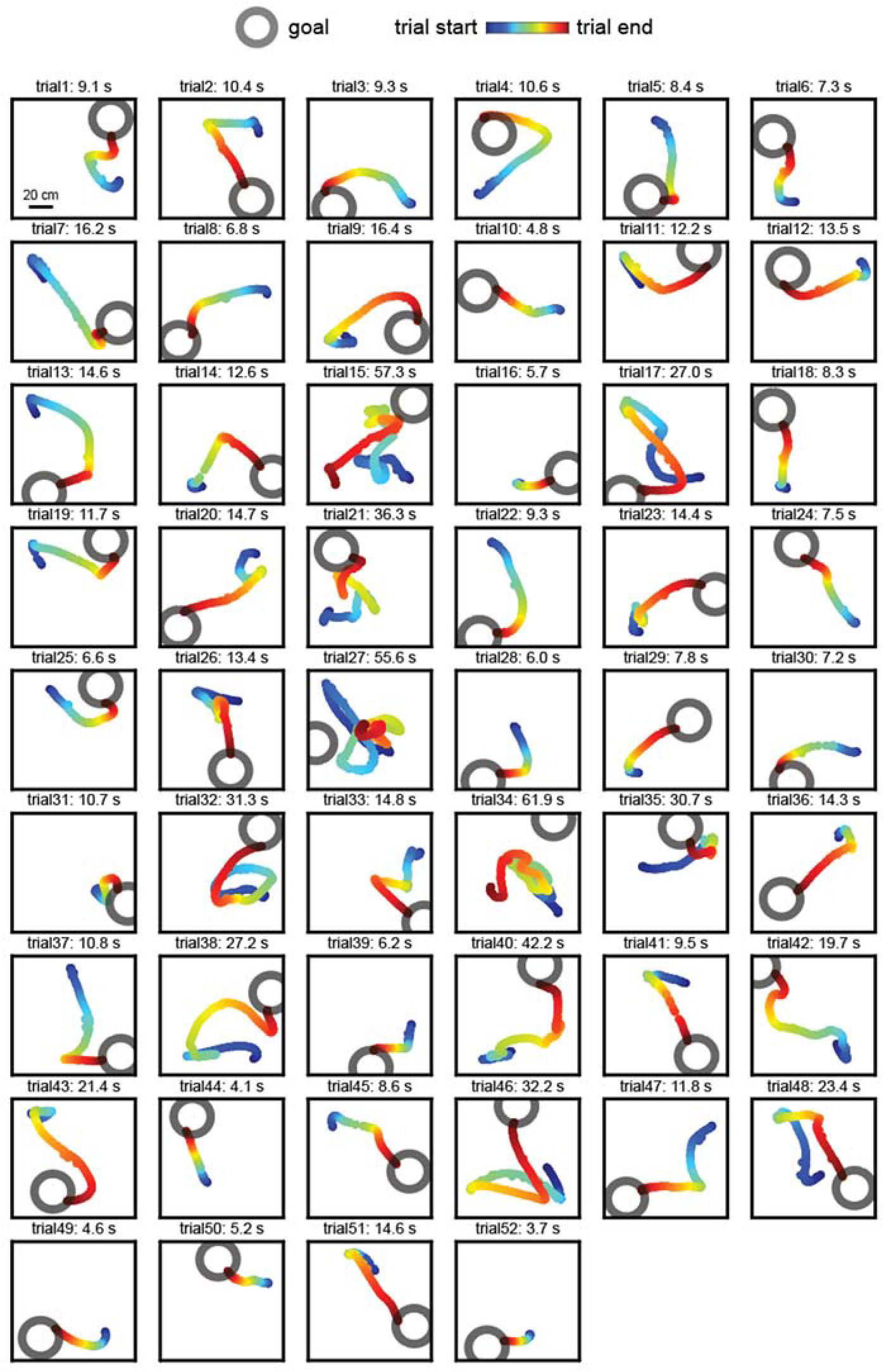
All trials of the Jumper BMI experiment for rat 2.

**Figure S5.**
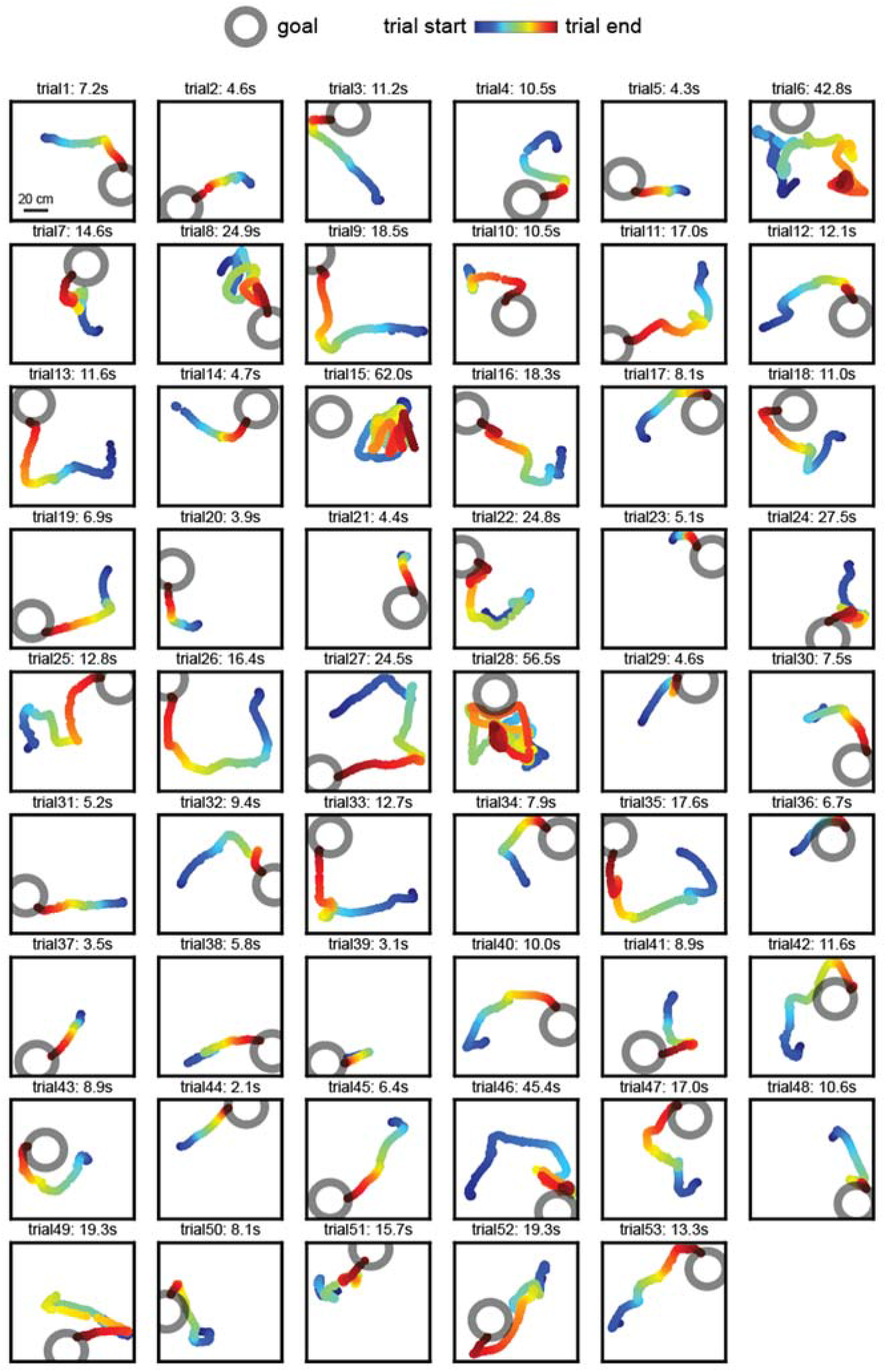
All trials of the Jumper BMI experiment for rat 3.

**Figure S6.**
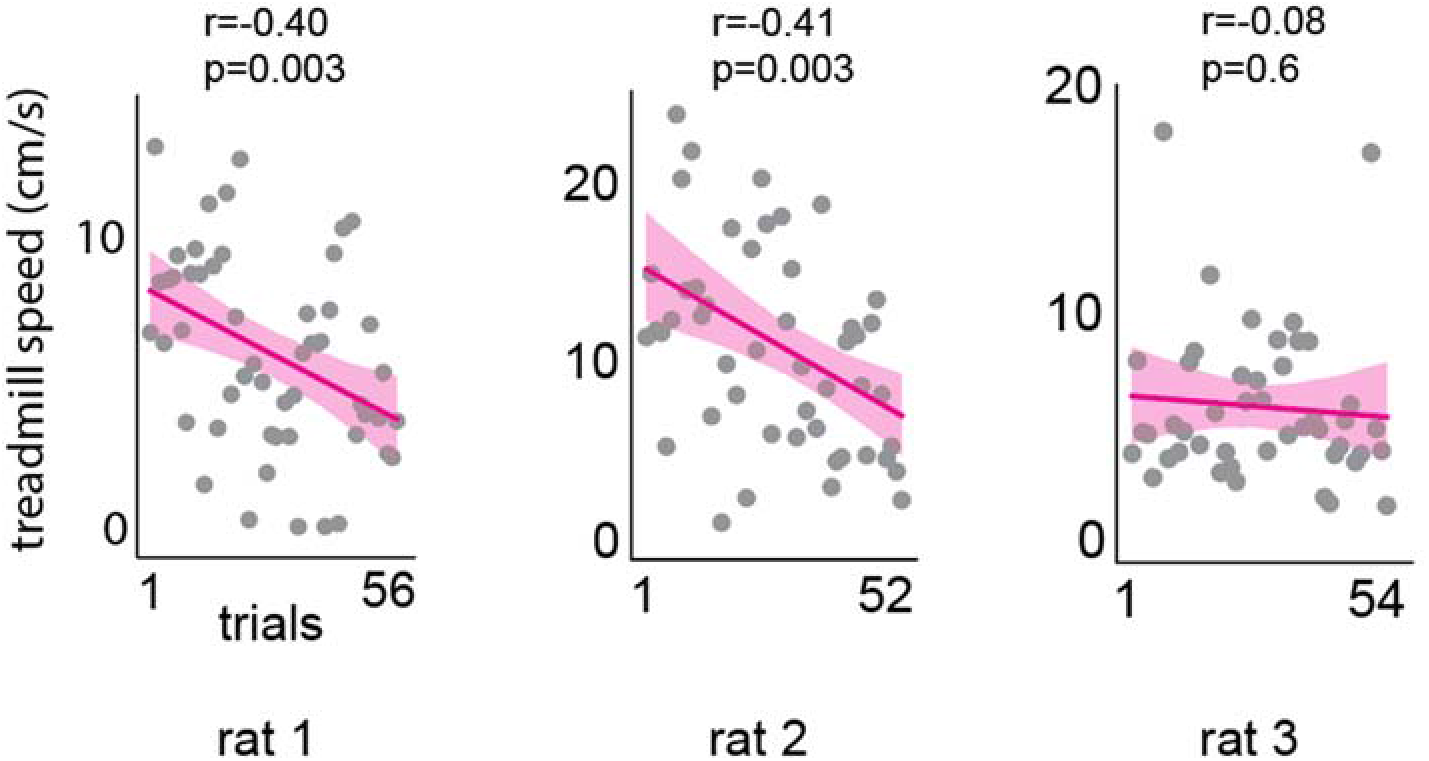
Mean treadmill speed during each Jumper trial and linear regression. Here and elsewhere all CIs are 95% CIs. For rats 1 and 2, treadmill movement decreased as the session progressed, while for rat 3 movement at the beginning was lower and remained at that level.

**Figure S7.**
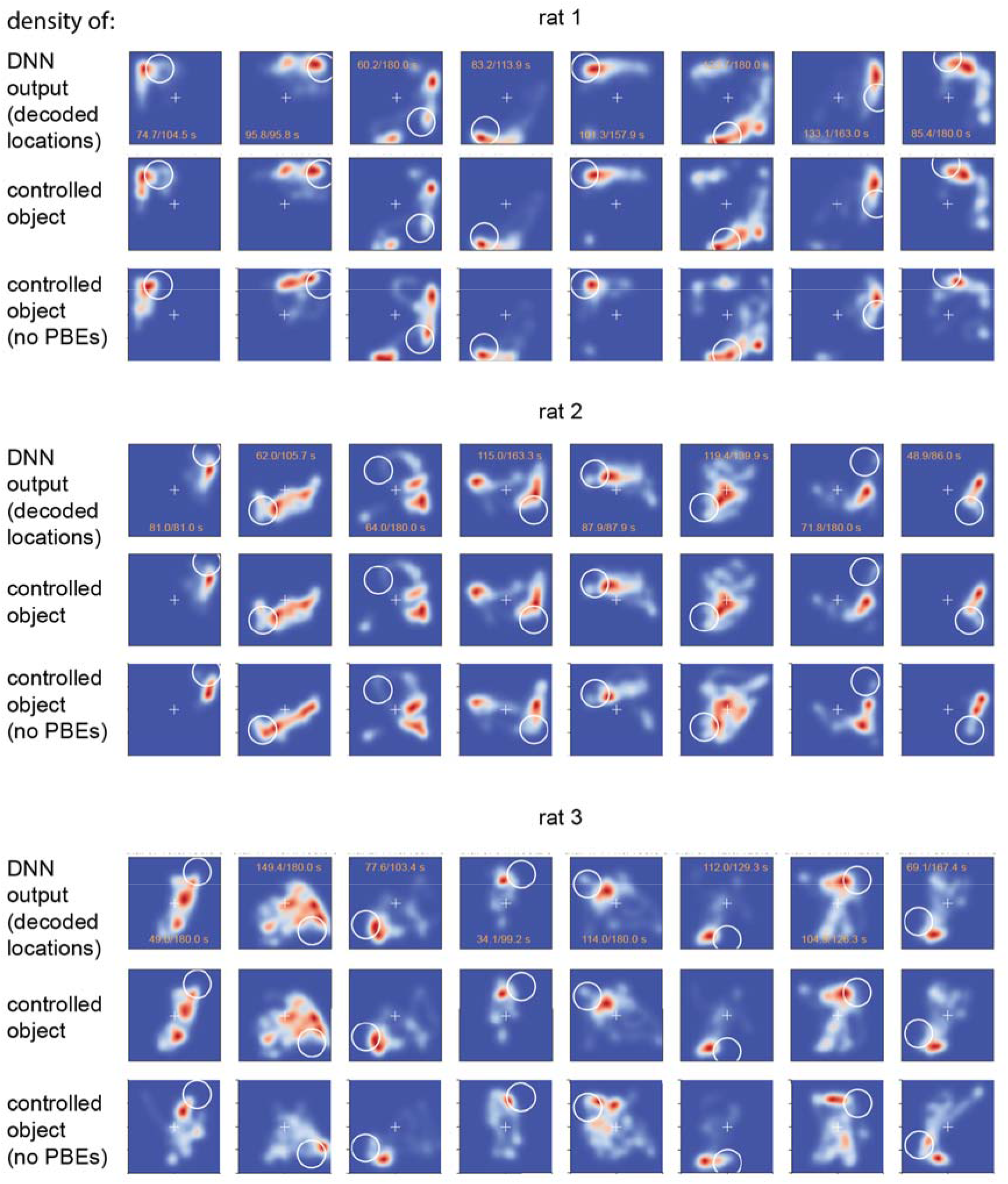
Distribution of real-time decoded location (i.e., DNN output), real-time controlled object location, and controlled object location after excluding activity during population burst events (PBEs) during the Jedi task. For each animal, the three rows represent the same 8 consecutive trials. Top row per animal is the same as shown in Figure 3A. Numbers in each panel indicate duration of trial excluding periods when animal’s angular velocity was >12°/s (a measure of task disengagement) (numerator) and total duration of trial (denominator), in seconds. In the bottom row per animal, spiking activity during detected PBEs was eliminated, then the location decoder was applied to the remaining data post-hoc. Comparison with the rows above shows that goal-directed control of the location of a remote object does not depend on this brief hippocampal population burst activity. Overall, the distributions are very similar across all conditions (rows).

**Figure S8.**
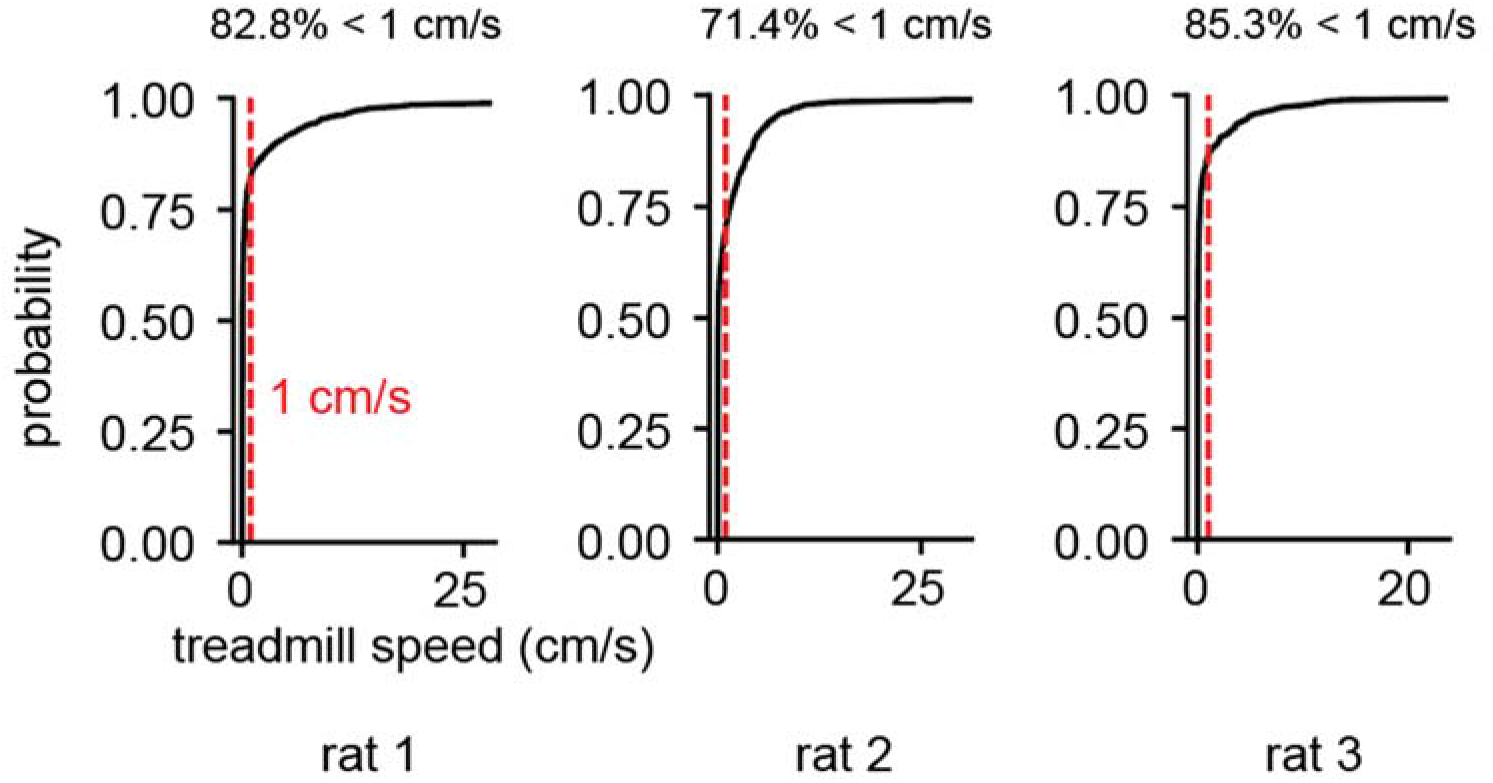
Treadmill speed during the Jedi task. CDF of the treadmill speed during periods across all trials when the angular velocity was <12°/s, i.e., the same periods the decoded location distribution is shown for in Figure 3 and Figure S7) and the fraction of samples when the treadmill speed was <1 cm/s, showing the animal was stationary most of time.

## Notes

### Competing Interest Statement

The authors have declared no competing interest.

